# Building the Next Generation Workforce: Why We Need Science Policy Training at the Undergraduate Level

**DOI:** 10.1101/2023.04.15.537026

**Authors:** Gwendolyn Bogard, Erin Saybolt, Moraima Castro-Faix, Adriana Bankston

**Author notes:** Authors contributed equally.

## Abstract

Diverse pathways into the science policy workforce are necessary for providing opportunities to students at all levels to engage in the field and to bring their talents to the future of policymaking. Both classroom and experiential training opportunities are essential to achieving this goal. In this publication, we assessed the current landscape of science policy training for undergraduate students in the United States by conducting a keyword search for experiential career training opportunities at varying levels of education. We also reviewed existing science policy publications for undergraduate opportunities, an area that has been largely unexplored to date. From these assessments, we found that most experiential training opportunities are geared towards Science, Technology, Engineering and Math (STEM) students in PhDs program or recent PhD graduates, followed by Masters-level students or graduates, and fewer opportunities exist for undergraduate students and Bachelor-degree holders. We then conducted focus group-style interviews with early career scientists involved in science policy, including undergraduate students, to discuss their experiences and ideas for change. Based on our findings, we recommend that universities, organizations, and funding agencies expand upon their existing resources for curriculum, academic advising, training opportunities, and funding, and dedicate resources toward raising awareness and creating additional opportunities in science policy at the undergraduate level. These steps can help create pathways for undergraduate students to enter and contribute to the science policy workforce.

## Introduction

### Current Landscape of Early Career Science Policy Training in the United States

Science policy is defined as the “set of federal rules, regulations, methods, practices, guidelines under which scientific research is conducted.”^1^ The field continues to expand as today’s societal challenges reveal the need for unique interdisciplinary approaches, as well as for showcasing the value of policymaking to incorporate evidence into the decision-making process. Science policy thereby has significant impacts for societal change at all levels of government, from the state and local landscape to the national and global stage.

Additionally, as policy change impacts every individual in our society, it is imperative to ground the policymaking process in diverse and inclusive viewpoints and backgrounds. The science policy workforce - which we define as professionals who transitioned into a science policy career from a variety of backgrounds and career stages - is not exempt from this principle, when it comes to researchers interested in Science, Technology, Engineering & Mathematics (STEM) fields applying their scientific knowledge and background to the policymaking process. Without voices that represent the wide-ranging needs of stakeholders invested in societal change, the lack of participation in the science policy workforce constrains the resulting policymaking process to benefit a narrower subset of society and only allows a small segment of the population to contribute. Evidence-based science policy is thereby critical for maintaining the competitiveness of the research enterprise in the United States, and the science policy workforce needs to reflect the people it serves.

The availability of diverse pathways for researchers interested in entering the science policy workforce is necessary for providing opportunities for STEM students at all levels to engage in the field and to bring their talents to the future of policymaking, including undergraduate and graduate students. Currently, science policy training across universities (for those in STEM and other disciplines) is highly variable, with few comprehensive training programs existing at the PhD level on a national scale. Given many of these opportunities are geared towards STEM students in a PhD program or recent PhD graduates, only a limited number of STEM undergraduate students across the country have access to science policy courses, which places this important group of students with an interest in science policy careers at a disadvantage for building their future careers.

Currently existing science policy training in universities—albeit still limited—is largely focused on STEM graduate students or recent PhD graduates in the form of elective courses, and some opportunities for Master’s programs in science policy. A few examples of Master’s programs include Georgetown University Master’s in Biomedical Science Policy & Advocacy,^2^ University of Wisconsin-Madison Neuroscience and Public Policy Program,^3^ and Massachusetts Institute of Technology Master’s in Technology and Policy,^4^ among others. Likewise, a few certificate programs in science policy across the country include: Science and Technology in Public Policy (STPP) graduate certificate program at University of Michigan,^5^ UC Irvine Certificate Program in Science Policy and Advocacy (as part of GPS-STEM),^6^ and UCR Science to Policy Certificate Program,^7^ in addition to a number of science policy fellowship programs.^8^ However, in many cases, eligibility for these programs presupposes that one must pursue PhD training in STEM in order to enter the science policy workforce through these opportunities.

Not all students across the country may be able to pursue or be interested in going the PhD route in STEM to achieve their career goals in science policy, including undergraduate students who may be seeking various other pathways to enter the field. This brings up the broader question of the currently existing landscape of science policy training, and the fact that novel pathways into the field should be created that facilitate the entry of undergraduate students into science policy careers through pathways different from a graduate degree. This expansion would help develop a broader framework for how undergraduate versus graduate students can enter the science policy workforce and contribute to societal change at different stages in their careers and through different mechanisms during their training.

At the undergraduate level, currently relatively few science policy opportunities exist, thereby limiting potential pathways for tomorrow’s leaders to enter and thrive in science policy careers (**Figure S1**). While some organizations allow undergraduate members to participate in science policy opportunities, the relatively small number of science policy opportunities limits talented students from contributing to the science policy workforce. Consequently, there is also a general lack of understanding and awareness of science policy among undergraduate students, some of which is outlined in this publication. When examining science policy training specifically for undergraduate students, the field lacks a clear and consistent pathway for them to explore career options in science policy, thereby blocking pathways for professionals with varying levels of education and professional interests who could contribute to the field. Recently, a few such examples have come to light, with more research needed to identify undergraduate opportunities in science policy in the classroom across the country.

As an example, in fall 2020, the course Interdisciplinary Applications of Biology^9^ became a fruitful environment for a collaboration between Georgia Gwinnett College and the *Journal of Science Policy and Governance (JSPG)*,^10^ an internationally recognized non-profit organization and peer-reviewed publication dedicated to empowering early career scientists, engineers, and policy professionals in international science policy debate. In this manner, science policy education was offered to undergraduate students in the course using the journal’s publications as a training tool to learn about science policy topics, as well as the process of writing a science policy paper and discussing these ideas in larger forums. While limited, this model has proven successful and could be further expanded to train undergraduate students across the country in science policy by collaborating with journals or other organizations that bring in science policy training opportunities and additional expertise into the classroom.

### Publication Summary and Recommendations

In this publication, we performed several initial analyses to assess the current landscape of science policy training for undergraduate students. We examined science policy opportunities that both current undergraduate students and bachelor’s degree-holders in STEM fields could apply to using currently available databases, conducted focus group-style workshops with students including undergraduate students, and performed a keyword analysis on science policy publications at different career stages including “undergraduate.” This pilot study revealed a lack of training opportunities in the science policy at the undergraduate level in STEM, highlighted the gaps in science policy education and workforce development, and laid out recommendations for filling these gaps. Our recommendations for change across several stakeholders to support undergraduate students in science policy focus on increasing awareness for training opportunities in universities, expanding current and developing additional training opportunities by various organizations, and incentivizing change through in the system specific funding mechanisms to support this goal.

### Undergraduate Science Policy Training Efforts at the STEM Advocacy Institute

The STEM Advocacy Institute (SAi)^11^ is an incubator that seeks to enable and accelerate the building of new tools and programs for expanding pathways of access between science and society. As part of SAi, The Bankston Lab^12^ is looking to build out the organization’s policy branch and develop innovative ways through which science policy can be a bridge between science and society. Given SAi’s strong focus on increasing participation of all critical stakeholders in strengthening the connections between science and society, focusing on training opportunities for undergraduate students and Bachelor’s degree-holders in STEM is an important segment of the workforce to be addressed.

Given the lack of training opportunities for undergraduate students with STEM or those in other majors interested in science policy, the Bankston Lab currently focuses on this group. To cultivate the next generation workforce in science policy, the Bankston Lab aims to perform research studies and develop training programs for undergraduate students, including for those in underrepresented backgrounds. These efforts are needed to understand the state of play in the field and the level of currently available opportunities for undergraduate students, with the goal of incentivizing the development of science policy training programs within universities and other organizations, as well as funding mechanisms to support science policy education. As part of the larger science policy ecosystem, through The Bankston Lab, SAi plans to create resources for undergraduate students to help them progress into the field and contribute to the science policy workforce.

## Methods

To better understand the needs of undergraduate students in science policy, we performed several pilot research studies and organized several workshops upon which we developed stakeholder recommendations.

### Review of Training Opportunities

To learn more about science policy training opportunities available to undergraduate students in STEM and other fields, we conducted a search for active experiential career training opportunities including fellowships and internships which undergraduates in the U.S. may be eligible for using a crowd-funded document^13^ and cross-referencing with the key words “science policy” to identify relevant opportunities, in addition to <u>NSPN’s resource center</u> and additional manual searches. Through this analysis, we also determined the education or career stage necessary to participate in each opportunity (**Figure S1**). The original <u>crowd-funded document</u> included other opportunities that are currently inactive, have specific citizenship requirements or are found in other categories, which we did not include in this figure.

### Group Interview Workshops

In a series of virtual events held in spring 2022, we spoke directly to trainees who are currently on the science policy career track to understand the challenges and opportunities they would like to have to pursue a career in the field. Given the nature of these workshops and the organizations that held them, although we did not quantify the field of study, it is safe to assume that many of them were in STEM fields and these data are also relevant to student researchers who may want to transition into policy. The data were obtained from two separate pilot workshops in which participants - majority early-stage graduate students and some undergraduates - were asked to respond to two prompts. One was to “identify problems with current pathways into the science policy workforce,” and the second was to “envision solutions that would address these problems with the science policy career pathway” to better support undergraduate students interested in science policy. In both workshops, we also held focus group-style discussions and identified key themes from the responses. There were in total 15 participants in both workshops. Responses from workshop participants were recorded, coded, and classified into several categories by moderators, and used to observe patterns using the Chi methodology.^14^

### Research of Publications for Undergraduate Policy Opportunities

Through a keyword analysis on Google Scholar, we analyzed the number of science policy publications at different career stages to determine the level by which publications could serve as opportunities to provide training in science policy as well as the potential level of exposure to the field. We searched scholarly publications in Google Scholar (using the “all time” function) for several combinations of keywords: 1) “science policy” alone as a term; 2) “undergraduate” AND “science policy”; 3) “graduate” AND “science policy” (in relation to graduate school); or 4) “masters” AND “science policy,” and quantified the results using the number of publications in the “science policy” alone category as a reference point.

## Results

### Review of Training Opportunities

In total, we found 79 active science policy training opportunities within our parameters (**Figure S1**). Although the list is not comprehensive, we believe it closely reflects the type and scope of opportunities which undergraduate students interested in science policy would encounter through general online searches. The resulting figure included active opportunities at the undergraduate level (red), or graduate level (Master’s or PhD) (black) based on the eligibility requirement for the specific science policy opportunity in the United States. The analysis included the following career stages: undergraduate student (current student or recent Bachelor’s graduate, column E), Master’s student or graduate (column F), and PhD requirement for the opportunity (column G).

From our list, the following organizations have opportunities that are currently active for current undergraduate students or recent Bachelor’s graduates (red text in **Figure S1**), which include: universities (Princeton University); non-profit organizations and scientific societies (American Association for the Advancement of Science; American Geophysical Union; American Mathematical Society; National Academies of Sciences, Engineering, and Medicine; Institute for Defense Analyses; Thriving Earth Exchange; Congressional Hispanic Caucus) and government agencies (Oak Ridge Institute for Science and Education; Department of Energy Office of Science and Energy Efficiency and Renewable Energy).

Based on our web search parameters and quantification of training options **Figure S1**, most active training opportunities in science policy are open to graduate students in PhD programs or PhD graduates in STEM (95%), whereas fewer opportunities exist for graduate students in Master’s programs or Master’s graduates (58%), and even fewer for undergraduate students or Bachelor’s degree graduates (16%) (**Figure 1**).

**Figure 1:**
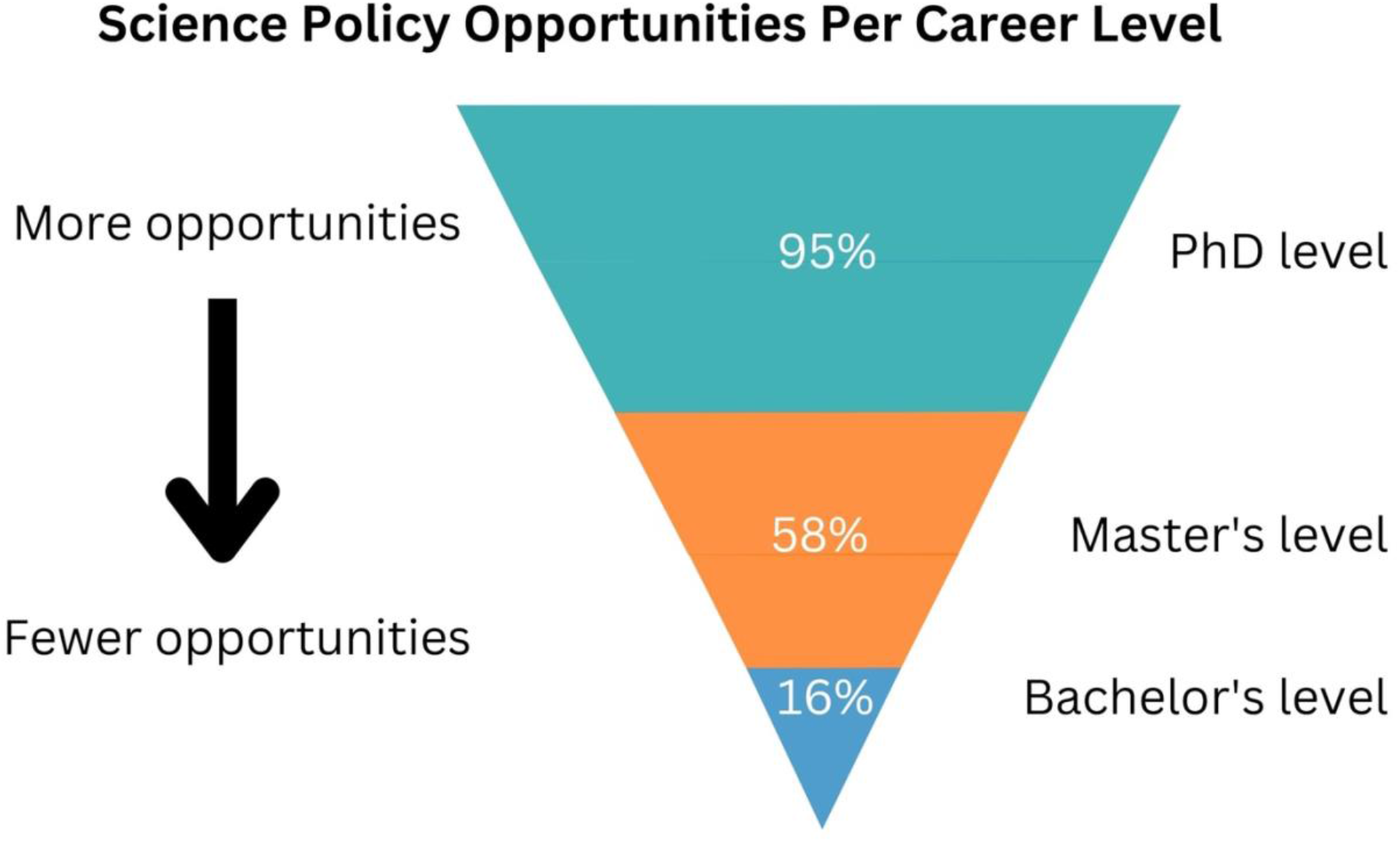
Science Policy Opportunities Per Career Level. Relative percentage of active science policy opportunities available for current students or graduates from PhD, Master’s, and Bachelor’s level programs based on **Figure S1**.

We then looked specifically at fellowships from our data, undergraduate students or Bachelor’s graduates were only eligible for 12.7% of fellowships out of the total number of opportunities available to all levels. In addition, a very limited number of entities host these opportunities across sectors, including member societies, universities, government agencies and nonprofits (red text in **Figure S1**). In comparison, Master students or Master’s graduates were eligible for 54.4% of fellowships, whereas PhD students or PhD graduates were eligible for 89.8% of fellowships out of the total number of opportunities available to all levels. These efforts showed a very small number of science policy fellowship opportunities at the undergraduate level compared to Master’s and PhD levels. When we looked at the internship availability from our data, this was very low (less than 5%) for all three educational levels, which indicates the broader need for such opportunities across the training landscape to expand the science policy workforce (**Figure 2**).

**Figure 2:**
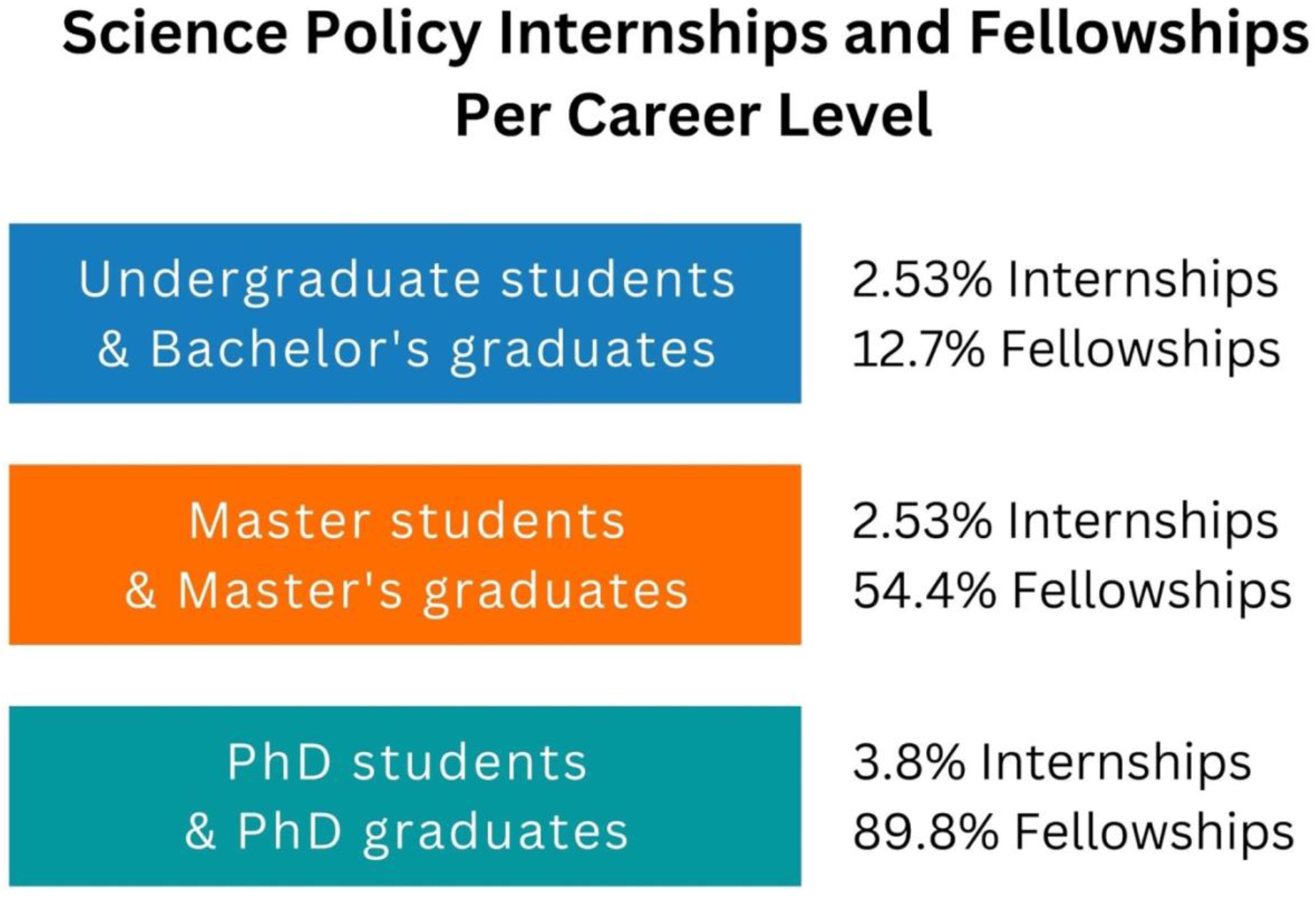
Science Policy Internships and Fellowships Per Career Level. Relative percentage of active internship and fellowship opportunities available for current students or graduates from the Bachelor’s, Master’s, and PhD level programs to enter the science policy workforce based on **Figure S1**.

### Group Interview Workshops

In addition to researching existing science policy opportunities, we interviewed graduate and undergraduate students in a workshop format to inquire about problems that undergraduate students may encounter when entering science policy, and asked participants to offer solutions.

When prompted to discuss problems that undergraduate students may encounter with entering science policy careers (**Figure 3**), workshop participants brought up several issues. In these workshops, 40% of participants reported the lack of exposure for undergraduate students to science policy opportunities. When asked why this may be the case, 28% of participants cited that undergraduate training in STEM is mainly focused on preparing students for academic or medical careers, which may lead to them not considering other career options. Workshop participants indicated that the perception for why a science policy career path may be unavailable to undergraduate students is because they are either unaware of the opportunities available in science policy training at the local level (21%), or the students may be unable to participate in fellowships or other science policy opportunities available to Master’s or PhD students (11%). These data are consistent with our prior findings, showing that science policy opportunities are much more limited at the undergraduate and Bachelor’s levels as opposed to other career stages when it comes to STEM disciplines.

**Figure 3:**
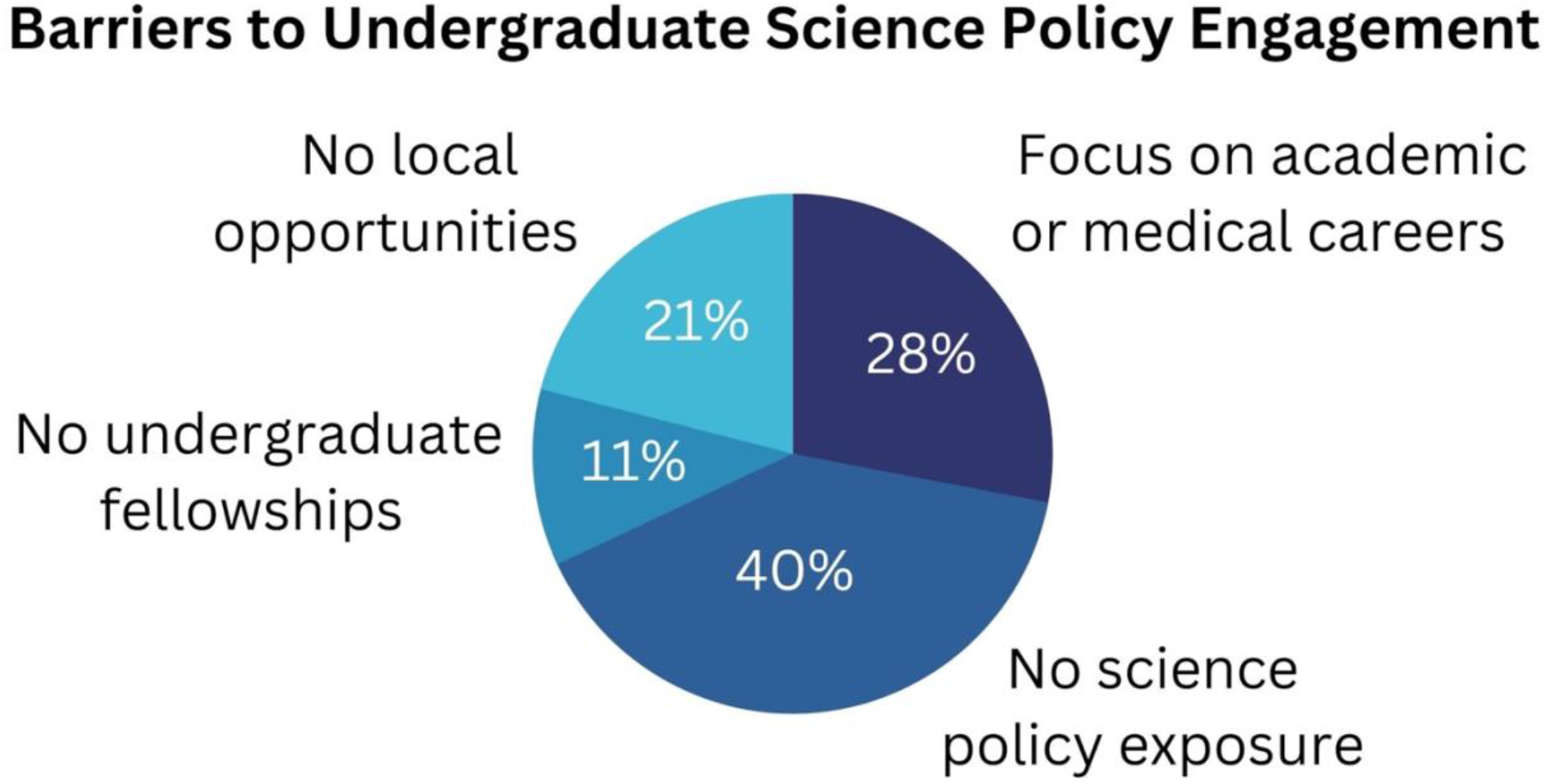
Barriers to Undergraduate Science Policy Engagement. Problems identified with current pathways into the science policy workforce for undergraduate students from the workshops. A table of quotes for each category (problem) can be found in **Appendix A**.

When prompted to discuss possible solutions to the lack of accessibility of policy careers to undergraduate students (**Figure 4**), workshop participants provided several solutions. A total of 40% of workshop participants indicated that undergraduate students may benefit from science policy courses to provide them with exposure to the field, whereas 20% of participants suggested that science policy post-baccalaureate (post-bac) opportunities in science policy could give undergraduate students an understanding of what it might entail to engage in these activities without pursuing additional degrees to enter the field. Other possible solutions to engaging undergraduate students in science policy from workshop participants included science policy career counseling (20%) or the development of a policy liaison position (20%) at a given school, which could facilitate participation of undergraduate students in these science policy initiatives to expand their knowledge of the field.

**Figure 4:**
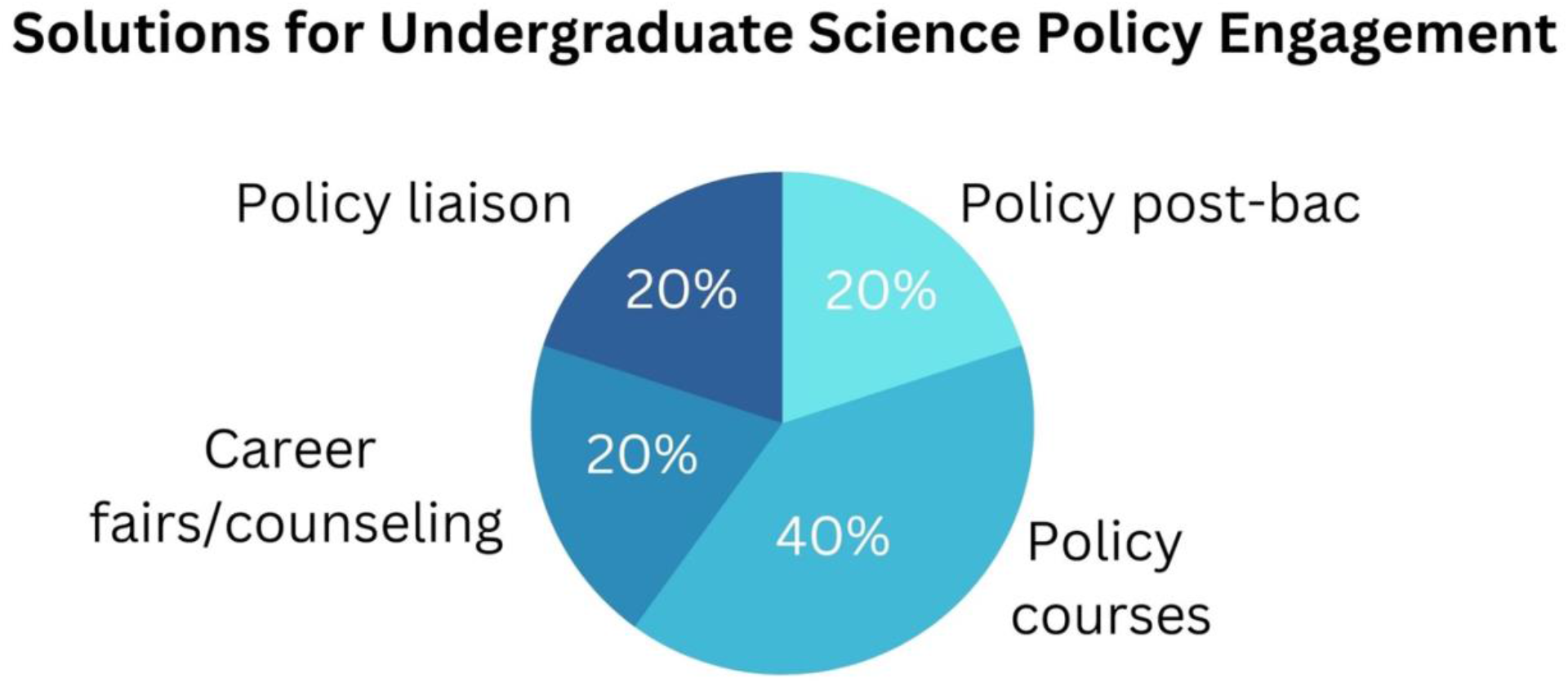
Solutions for Undergraduate Science Policy Engagement. Proposed solutions to problems (identified in Figure 3) of engaging undergraduate students in science policy from the workshops. A table of quotes for each category (solution) can be found in **Appendix B**.

### Research of Publications for Undergraduate Policy Opportunities

To examine science policy opportunities through publications, we searched science policy articles that relate to different career stages. In contrast to searching scholarly publications in Google Scholar for the word “science policy,” which yielded a large number of publications (termed 100% for reference), relatively to this number, combinations of the word “graduate” AND “science policy” led to a percentage of 3.3%, whereas the combination of search terms “masters” AND “science policy” led to a percentage of 0.7%, and the combination of search terms “undergraduate” AND “science policy” was only 1.1% (**Table 1**). In this case, the masters and undergraduate categories were close to each other, indicating that both career stages could use more science policy opportunities to be provided and written about to increase exposure, as opposed to the available and visible options at the PhD level.

**Table 1:** Science Policy Term Search in Publications. Keyword search of scholarly publications with various terms in combination with the term “science policy” (Google Scholar). The data in the table refer to the ratio of publications with the combination of keywords in each category as relative to the “science policy” publication number alone, which is 2,100,000 number in the “science policy” category (*) as a percentage.

| **Combination of keywords** | **Number of publications with given keywords\*** | **Percentage publications with given keywords relative to “science policy”** |
| --- | --- | --- |
| “science policy” | 2,100,000 | 100% |
| “graduate” AND “science policy” | 69,700 | 3.3% |
| “masters” AND “science policy” | 15,400 | 0.7% |
| “undergraduate” AND “science policy” | 22,200 | 1.1% |

## Discussion

Policy solutions to issues involving science and society rely on participation from individuals at all career levels who can share a variety of ideas and perspectives to ensure future solutions maximize the good for society. Science policy is a critical piece of this process and similarly requires opportunities early in training, including at the undergraduate level, to provide opportunities for engagement in the science policy workforce.

Our research showed that science policy opportunities for undergraduate students are limited in number, and that these decrease when moving from PhD to Master’s level programs or degrees and finally to the undergraduate level (**Figure 1**). For students who do not wish to pursue these degrees or are seeking to enter science policy through other means, limited opportunities in terms of fellowships (currently the most popular means of transitioning into the field) and internships (**Figure 2**) can lead to the exclusion of this population from the science policy workforce at the national and international levels, as well as in the state and local arenas.

Our workshops shed light onto several problems that undergraduate students may experience that precludes them entering the field of science policy (**Figure 3**), such as the lack of professional opportunities that Master’s and PhD-level graduates are eligible for. In addition, the reduced exposure of undergraduate students to knowledge of existing programs they could benefit from will also disadvantage them. Solutions proposed in the workshop include additional training, as well as talking with individuals who can offer educational assistance such as career counseling (**Figure 4**).

Our analysis of keywords in Google scholar for science policy publications using “science policy” with additional terms and career stages similarly revealed the limited number of opportunities that may be found for each of these career stages as opposed to more advanced ones in the list (**Table 1**). Whereas more results came up with the “graduate” as a term (in relation to graduate school), the search yielded a relatively small percentage of results for all career stages noted, which were even lower at the Master’s and undergraduate levels. These data indicate a broader need for more available information on these opportunities for students at all levels, including for undergraduate students to find and take advantage of.

### Study Limitations

Given this was a small pilot study to assess the landscape and current needs for undergraduate students in science policy in STEM and other areas, our analysis was limited in scope. First, the science policy lists upon which we based our search, whereas comprehensive, likely did not contain all the possible opportunities that exist for undergraduate students interested in the field. Second, the number of workshop participants represents a limited subset of views based on what we were able to gather from a small group. Third, the Google Scholar keyword analysis was limited by keywords and online resources that have been created at the time when the search was done. Science policy is a broad field and as such, opportunities may have been missed if using other descriptors for the term science policy, which would be excluded from the search.

Nevertheless, with this study we aimed to gather some preliminary results to map a path forward for follow up studies to build on in terms of undergraduate opportunities in science policy. Therefore, we believe this pilot study is a valuable resource for the science policy community and a way to learn more about an area that has been examined very little to date when it comes to the undergraduate level and potential pathways into the science policy workforce that may not have been previously considered or explored by students in STEM and other disciplines. We hope this work will lead to more analysis of this landscape in the future. In general, our data show that significant gaps exist in science policy training at the undergraduate level for several reasons, and the study indicates a few areas where improvement could be made in the field. We believe that further research is warranted to increase the sample size with additional data and views represented by the next generation based on this study, and that additional studies will provide a full picture of the needs in this space.

### Future Studies and Recommendations

In a broad sense, utility could be found in interviewing students in undergraduate programs about their understanding of the science policy field and what resources may be most useful for them in a larger scale study driven by this initial analysis. More specifically, this work could lead to the development of more science policy opportunities by various stakeholders, especially in the form of internships and potentially specific fellowships geared specifically towards undergraduate students entering the field. Other options include expanding eligibility of current opportunities (such as science policy fellowships) to undergraduate students or Bachelor’s-level graduates or creating other specific avenues for them to engage in science policy similarly to those that already exist for Master’s and PhD-level graduate students.

Given that science policy fellowships are one of the most popular avenues by which graduate students have transitioned into the field, further discussion in their expansion to undergraduate students is warranted without having to pursue PhD degrees. Even so, several fellowships require living in the nation’s capital to engage in their programs, which may be another barrier for students unable to make the move and take advantage of this stepping stone. While some state fellowships exist,^15^ many are currently limited across the country in terms of geography and likely also limit eligibility to graduate students. In the case of undergraduate students, expansion of federal or state level fellowships would need to occur broadly by creating opportunities across the country.

The lack of training opportunities for undergraduate students in science policy is another major barrier to engagement in the field. Indeed, as evidenced by this study and others, currently, universities across the country could improve their teaching curriculum to include science policy fundamentals. Anecdotal evidence suggests that most students learn about science policy opportunities when they are already in a graduate program, and on their way to graduating with a Master’s or PhD in STEM or other fields. While these are very valuable pathways for transitioning into the field, not every undergraduate student has the means to apply for and attend such a program to pursue this career path. In general, barriers that exist for undergraduate students in science policy training can be addressed through even more information-sharing efforts than the currently existing resources (some of which were also used in this study) and additional opportunities for undergraduate students in the form of internships and fellowships that will need to be developed specifically for or eligibility could be extended to this population from various stakeholders.

To an extent, the culture change that needs to occur within universities allowing for science policy training to be integrated into the curriculum will need to include a more formal exposure of undergraduate students to the science policy career path. This will require changes at all levels such as: 1) making academic advisors more aware of opportunities in the field for their undergraduate students; 2) working with academic staff to develop courses, seminars and mentoring opportunities to expose undergraduate students to science policy, training and workforce development; and 3) identifying ways to expand undergraduate student eligibility for current opportunities which Master’s and PhD level students already take advantage of.

In addition to fellowships, undergraduate students could apply for science policy internships, however currently these opportunities are fairly limited as well and may be more difficult to find. Such internships could be valuable opportunities that universities can advertise more to expose undergraduate students to a variety of placements, including Capitol Hill, government offices or non-profits as a few examples. Due to the lack of centralized resources for identifying science policy internships, many students individually search for them, and may also not have exposure to others who have previously enrolled in the program to get an idea of the day to day.

It is also worth noting that, while expanding current PhD-level science policy fellowships and internships to undergraduates is a potential option, undergraduate students often have much less expertise in science policy. Therefore, the expansion of current opportunities or the development of new opportunities for undergraduates in science policy should be exploration-oriented, providing students with a wide variety of information on science policy opportunities and career tracks. This does not need to be limited to training opportunities outside of the university, but rather integrating science policy training offered by organizations and non-profits into the undergraduate curriculum is an opportunity to increase awareness of these options. For students who may not want to pursue a Masters or PhD-level degrees, this is a desirable option for undergraduate students interested in science policy, offering an alternative path for entering the field.

### Diversity in Undergraduate Science Policy Opportunities

One aspect we did not delve into deeply in this publication, but which should be kept in mind moving forward in these studies, is that undergraduate students from underrepresented backgrounds are often at a disadvantage for opportunities in science policy, and working to change this can further enrich the field using a large variety of perspectives. Barriers to inclusion of undergraduate students from various backgrounds in science policy, which we also aim to address at SAi, constitute the limited number and type of opportunities currently existing which these populations may be aware of or be able to take advantage of due to several reasons,^16^ and this creates a less diverse field of science policy professionals.

In addition, undergraduate opportunities and support that exist in science policy within universities or from other stakeholders are rarely housed or accessible at institutions which serve low-income and underrepresented populations, which also places a large proportion of talented undergraduate students at a disadvantage for educational and career opportunities in the field. Future work should focus on creating science policy options in different types of research universities including HBCUs and MSIs, as well as become as a priority by other stakeholders^17^ through internships that can also help low-income students participate in these opportunities. In addition to internships, other ways for undergraduate students from underrepresented communities to gain exposure to science policy include committees^18^ and providing additional academic advising support, as well as financial support if needed during these programs. These efforts should be undertaken using intentional recruitment strategies that allow for underrepresented students to apply for these opportunities and participate in training and career development in science policy in relation to developing the future workforce.^19^

Finally, although we did not address it in this study, science policy opportunities for international students, including undergraduate students interested in science policy careers, are also limited due to several factors. Offering additional assistance and developing resources particularly for undergraduate students who are foreign-born and wish to pursue science policy opportunities in the United States which have been outlined in this publication should be an area of focus for various stakeholders moving forward to build future leaders in the field.

## Recommendations for Change

To bolster representation of undergraduate students in science policy training and the future workforce, we propose three broad recommendations that will strengthen future national and local efforts (**Table 2**). These recommendations are detailed below as a broader call to action for universities, organizations, and funding agencies, in terms of offering training and mentoring, as well as additional resources and partnerships that could help expand opportunities, and grant mechanisms where science policy opportunities can be considered.

**Table 2:** Stakeholder Recommendations for Undergraduate Science Policy Opportunities. Proposed recommendations to three stakeholders to address the lack of science policy opportunities for undergraduate students in the United States.

| **Universities** | **Organizations** | **Funding agencies** |
| --- | --- | --- |
| Science policy training in curriculum | Expand offerings to undergraduates | Expand policy sections related to research |
| Develop partnerships with existing programs | Create inclusive opportunities | Develop mentoring sections in grants |
| Advisors informed about science policy | Provide policy mentorship | Provide resources to connect science and society |

## Recommendations for universities

### Training and awareness of science policy opportunities and partnership development

Universities should include or enhance existing science policy training within their undergraduate curriculum, promote awareness of science policy as a career field, and develop partnerships with existing programs that already offer internships or other types of science policy training opportunities for undergraduate students.

- Professors should incorporate science policy topics into undergraduates STEM classes, with the help of existing resources;
- STEM departments should develop partnerships with organizations and government entities - such as Capitol Hill staff and agencies to offering experiential opportunities and communicate those to the undergraduate students;
- Academic advisors should be informed about science policy career paths and incorporate science policy opportunities into their career development and advising work to undergraduate students.

## Recommendations for organizations

### Expand existing professional science policy and educational opportunities for undergraduate students on science policy

Organizations, including scientific societies and nonprofits should expand their existing professional offerings in science policy and educational training in the field, such as career panels and more in-depth policy discussions, to include undergraduate students in the audience and showcase role models who have undertaken science policy training during their undergraduate years.

- Scientific societies and non-profits in the science policy space should create science policy programming aimed at undergraduate students;
- Current science policy opportunities at the undergraduate level offered by scientific societies and non-profits should be expanded across the country and include students from underrepresented backgrounds who are enrolled in STEM courses;
- Science policy professionals should provide mentorship to undergraduate students interested in science policy through programs developed by scientific societies and non-profits.

## Recommendations for funding agencies

### Research grants funding undergraduate students should include policy training

Funding agencies should expand sections of their existing research and training grants, or create new sections of these grants, that focus specifically on undergraduate student training and mentoring in science policy, including learning how to highlight the societal value of their research to policymakers.

- Research grants, or training grants designed for undergraduate students, should include science policy sections related to the research performed by undergraduates, which can incentivize the connection between science and society through their training;
- Offer mentoring opportunities in science policy to be undertaken by academic advisors or other administrative staff in various forms, and include outcomes of such mentoring in future grants that can used to showcase the development of skills highlighting the societal value of science to policymakers;
- Develop new research or training grant proposals or incorporate relevant sections into existing grants to help undergraduate students connect science and society; provide the necessary opportunities for this purpose, such as courses, seminars, trips to Capitol Hill, and exposure to science policy opportunities outside the university for experiential learning.

## Disclaimer

The views expressed in this publication represent the personal views of the authors and not the views of their employers.

## Figure Legends

**Figure S1: Science Policy Opportunities by Career Stage**. See the methods section for more details (<u>Supplementary Figure 1</u>).

## Tables

**Appendix A: Problems identified with undergraduate science policy training and quotes from workshop participants**. Workshop participants identified barriers preventing undergraduate students for engaging with science policy (see **Figure 3**).

| Problems identified | Participant quotes |
| --- | --- |
| Focus on academic or medical careers | "Traditional pathways (i.e. medical school) are highly encouraged at the undergraduate level." |
| Lack of exposure to science policy opportunities | "It is challenging to understand different career options (especially in basic science). Since there is not a lot of support from advisors, we need to seek help and do exploration on our own." |
| Science policy fellowships aren't accessible at the undergraduate level | "It can be intimidating for undergraduate students as seemingly fellowships are only available to PhD graduates." |
| Lack of local opportunities in science policy | "Local/state level sci pol is often forgotten - there are opportunities for diff levels of education here." |

**Appendix B: Solutions identified for undergraduate science policy training and quotes from workshop participants**. Workshop participants identified solutions that would enable undergraduate students to engage with science policy (see **Figure 4**).

| Proposed solutions | Participant quotes |
| --- | --- |
| Post-bac opportunity in science policy before graduate school | "Undergraduate students who recently graduated can explore science policy work before they go to graduate school through a post-bac opportunity." |
| Courses or seminars specific for undergraduates in science policy | "If you train students to have career development early, they may better understand science policy pathways." |
| Career fairs or counselors | "Being present at career fairs to connect with students about the science policy workforce." |
| Policy liaison | "Have students be liaisons between undergraduate students and science policy advocates." |

## Footnotes

1 *Beyond Sputnik*, accessed December 16, 2022, https://www.press.umich.edu/335760/beyond_sputnik.

2 “Biomedical Science Policy & Advocacy,” *Graduate School of Arts & Sciences* (blog), accessed November 8, 2022, https://grad.georgetown.edu/biomedical-science-and-advocacy/.

3 “Neuroscience and Public Policy Program – UW–Madison,” accessed November 8, 2022, https://npp.wisc.edu/.

4 “About TPP,” *Technology and Policy Program* (blog), accessed November 8, 2022, https://tpp.mit.edu/about/.

5 “STPP Graduate Certificate Program | Gerald R. Ford School of Public Policy,” accessed November 8, 2022, https://fordschool.umich.edu/stpp.

6 “UCI Graduate Professional Success for STEM PhDs and Postdocs,” accessed November 8, 2022, https://gps-stem.grad.uci.edu/.

7 “S2P Certificate,” Science to Policy, accessed November 8, 2022, https://sciencetopolicy.ucr.edu/s2p-certificate.s

8 “Policy Fellowship Database,” Genetics Society of America, accessed November 8, 2022, https://genetics-gsa.org/policy/policy-fellowship-database/.

9 “Cultivating the Policymaker Within: Using JSPG Publications in an Undergraduate Course,” Journal of Science Policy & Governance, accessed November 8, 2022, http://www.sciencepolicyjournal.org/4/post/2021/01/cultivating-the-policymaker-within.html.

10 “JSPG,” accessed December 16, 2022, https://www.sciencepolicyjournal.org/.

11 “STEM ADVOCACY INSTITUTE,” accessed November 8, 2022, https://stemadvocacy.org/.

12 “Bankston Lab - SAi,” accessed November 8, 2022, https://stemadvocacy.org/bankston-lab/.

13 “THE List of #SciPol Fellowships - Google Docs,” accessed November 8, 2022, https://docs.google.com/document/d/1-S407AQIu0cZI0HASAer1mBmZd57KXFMhKk79JnzBYk/edit.

14 Michelene T.H. Chi, “Quantifying Qualitative Analyses of Verbal Data: A Practical Guide,” *Journal of the Learning Sciences* 6, no. 3 (July 1, 1997): 271–315, https://doi.org/10.1207/s15327809jls0603_1.

15 “Developing Science and Technology Policy Fellowships in State Governments without Full-Time Legislatures,” Journal of Science Policy & Governance, accessed December 16, 2022, https://www.sciencepolicyjournal.org/article_1038126_jspg_16_01_04.html.

16 “Nearly a Quarter of Tenure-Track Faculty Have a Parent with a PhD | Nature Human Behaviour,” accessed November 15, 2022, https://www.nature.com/articles/s41562-022-01426-3.

17 “White House Initiative on Advancing Educational Equity, Excellence, and Economic Opportunity through Historically Black Colleges and Universities,” accessed November 15, 2022, https://sites.ed.gov/whhbcu/internshipsandfellowships/.

18 “Diverse Policy Committees Can Reach Underrepresented Groups,” *BFI (blog), accessed October 6,* 2022, https://bfi.uchicago.edu/working-paper/diverse-policy-committees-can-reach-underrepresented-groups/.

19 Meghna Sabharwal and Iris Geva-May, “Advancing Underrepresented Populations in the Public Sector: Approaches and Practices in the Instructional Pipeline,” *Journal of Public Affairs Education* 19, no. 4 (2013): 657–79.

